# Ciclopirox suppresses poxvirus replication by targeting iron metabolism

**DOI:** 10.1101/2025.07.24.666650

**Authors:** Anil Pant, Djamal Brahim Belhaouari, Lara Dsouza, D.M. Nirosh Udayanga, Zhengqiang Wang, Zhilong Yang

## Abstract

Poxviruses remain a significant global health concern, necessitating the development of novel antiviral strategies. Through high-throughput screening, we previously identified ciclopirox (CPX), an FDA-approved antifungal, as a hit that inhibits vaccinia virus (VACV) replication. Here, we further characterized its antiviral activity and mechanism of action using human primary fibroblasts. CPX significantly reduced VACV titers without reducing host cell viability, with an EC_50_ in the sub-micromolar range and a CC_50_ >500 μM. Rescue experiments demonstrated that CPX inhibits viral replication primarily through chelation of intracellular Fe^3+^ and, to a lesser extent, Fe^2+^, as evidenced by partial restoration of viral replication with ferric ammonium citrate supplementation. Furthermore, overexpression of the iron-dependent enzymes RRM2 and the VACV-encoded F4L reduced the inhibitory effect of CPX, indicating that these host and viral proteins are affected by CPX treatment. Moreover, CPX treatment also suppressed cowpox virus and monkeypox (mpox) virus replication *in vitro.* It also reduced VACV titers in *ex vivo* mouse lung tissue. These findings highlight host iron metabolism as a critical determinant of poxvirus replication and support repurposing CPX as a broad-spectrum orthopoxvirus antiviral candidate.

## Introduction

Poxviruses, including variola virus (the causative agent of smallpox) and monkeypox (mpox) virus (MPXV), pose significant threats to human and animal health. These threats may result from natural transmission, zoonotic spillover, or potential misuse as biological weapons. In addition to human diseases, several other poxviruses such as goatpox, sheeppox, and lumpy skin disease virus cause severe illness in livestock and contribute to economic losses and threats to wildlife. MPXV is endemic in Central and West Africa but has recently spread globally, with reported cases in more than 100 non-endemic countries. This rapid spread has raised concerns about the potential for another pandemic [1,2]. The combination of waning immunity to poxviruses, decreased global vaccination coverage, possibility of re-emergence from unsecured stocks [3], advances in synthetic biology [4], and increased international mobility underscores the urgency of developing new and safe antiviral strategies to counter smallpox, MPXV, and related poxviruses [5].

Two antiviral drugs, tecovirimat and brincidofovir, have been approved by the FDA for strategic stockpiling against smallpox [6,7]. However, brincidofovir has shown limited efficacy against MPXV [8,9], and is associated with serious adverse effects [10–12], and drug resistance in human cytomegaloviruses [13–15]. Although tecovirimat has promising activity against MPXV in some studies [16], the clinical efficacy data remain limited with no clear clinical benefits in humans against mpox [17–20]. Moreover, the risk of viral resistance against tecovirimat is a valid concern due to its low barrier to resistance [7]. Given the potential re-emergence of smallpox, the broad host range of poxviruses, and the emergence of drug-resistant or highly transmissible MPXV strains, there is a clear need to develop effective and accessible countermeasures.

The development of new antiviral drugs is a lengthy and expensive process, often requiring more than a decade of development and substantial financial investment. On the other hand, repurposing existing drugs with established safety and pharmacokinetic profiles offers a more rapid and cost-effective alternative. Ciclopirox (CPX) has been known as a broad spectrum antifungal since the early 1970s [21]. As an FDA-approved drug, CPX is currently used topically to treat conditions such as seborrheic dermatitis, dandruff, and nail fungal infections [22–25]. Recent studies have also reported potential applications of CPX in cancer and viral infections [26–33].

Our previous high-throughput screening of FDA-approved and bioactive compounds identified CPX as one of the top hits that efficiently suppressed vaccinia virus (VACV) replication [34]. Here, we characterized the antiviral activity against different poxviruses and molecular mechanism of CPX. We found that CPX efficiently suppresses the replication of VACV, cowpoxvirus (CPXV), and MPXV in primary human cells and *ex vivo* mouse lung tissues.

Mechanistic studies revealed that CPX inhibits VACV DNA replication and reduces post-replicative gene expression. CPX interferes with iron metabolism and, at least in part, suppresses the function of ribonucleotide reductase M2 (RRM2), a host enzyme required for the synthesis of deoxynucleotides. Together, these results identify CPX as a promising candidate for further development for managing poxvirus infections and provide insight into its antiviral mechanism of action.

## Materials and Methods

### Cells and Viruses

Primary human foreskin fibroblasts (HFFs) were generously provided by Dr. Nicholas Wallace (Kansas State University). HFFs and 293-FT cells were maintained in Dulbecco’s Modified Eagle Medium (DMEM; Fisher Scientific) supplemented with 10% fetal bovine serum (FBS; VWR), 2 mM L-glutamine (VWR), 100 U/mL penicillin, and 100 µg/mL streptomycin (VWR). BS-C-1 cells (ATCC CCL-26) were cultured in Eagle’s Minimum Essential Medium (EMEM; Fisher Scientific) with the same supplements. All cells were maintained at 37 °C in a humidified atmosphere with 5% CO₂.

The VACV Western Reserve (WR) strain (ATCC VR-1354) was amplified and purified by ultracentrifugation through a sucrose cushion as previously described. CPXV (strain Brighton Red) and MPXV-MA001 2022 isolate (GenBank: ON563414.3) were used in this study. Recombinant VACVs expressing *Gaussia* luciferase under early (vEGluc), intermediate (vIGluc), or late (vLGluc) viral promoters were described previously [35]. Virus preparation and infection protocols were performed as reported by previous publications [36].

### Virus Titration by Plaque Assay

VACV, CPXV and MPXV titers were determined by plaque assay as described previously [36]. BS-C-1 cells were seeded in 12-well plates and infected with serially diluted virus for one hour and then grown in culture medium containing 0.5% methylcellulose. After 48h, Cells were stained with 0.1% crystal violet for 15 min, followed by washing with water, and the number of plaques was counted.

### Chemicals

Cytarabine (AraC) was purchased from Sigma-Aldrich. Ciclopirox olamine (CPX) was purchased from TargetMol and was dissolved in DMSO for all experiments. Deferoxamine mesylate (DFO) was obtained from Ambeed. Ammonium ferric citrate (FeAmC) was purchased from MedChemExpress. Bathophenanthroline disulfonic acid (BPhen) and deferiprone (DFP) were purchased from Cayman Chemical. Magnesium sulfate (MgSO₄) was obtained from Fisher Scientific. Manganese chloride (MnCl₂), adenosine, uridine, guanosine, and cytidine were obtained from Sigma-Aldrich

### Cell Viability and CC₅₀ Determination

Cell viability was assessed using the trypan blue exclusion method and the CCK8 assay as described previously [37]. For trypan blue staining, cells were treated with DMSO or the indicated compounds at indicated concentrations and counted using a TC20 automated cell counter (Bio-Rad) after staining. For the CCK-8 assay, cells were seeded in 96-well plates and treated with DMSO or indicated compounds at indicated concentrations. After indicated time points, 10 µL of CCK-8 reagent was added to each well, followed by a 1 h incubation at 37 °C. The absorbance at 450 nm was then measured using a Cytation 5 imaging reader (BioTek). For the MTT assay, cells were seeded in 96-well plates and treated with DMSO or indicated compounds at various concentrations. After 24 or 48 h, 10 µL of MTT reagent (Cayman Chemical) was added to each well, followed by a 3 h incubation. Then, 100 µL of crystal dissolving solution was added, and the absorbance at 570 nm was measured using a Cytation 5 imaging reader (BioTek) after overnight incubation at 37 °C.

To determine the half-maximal cytotoxic concentration (CC₅₀), cells were treated with a range of compound concentrations, and viability was quantified using the MTT assay. CC₅₀ values were calculated in GraphPad Prism (version 10.2.3) using nonlinear regression analysis with the “log(inhibitor) vs. normalized response-variable slope” model. The concentration resulting in 50% reduction in viability relative to vehicle control was defined as the CC₅₀.

### Determination of EC₅₀

To determine the half-maximal effective concentration (EC₅₀), HFFs were seeded in 96-well plates and infected with vLGluc at an MOI of 0.01 in the presence of vehicle or indicated compounds at several concentration with serial dilutions. *Gaussia* luciferase activity was measured at 24 h post-infection (hpi) using a luminometer. EC₅₀ values were calculated by nonlinear regression analysis using the equation “log (inhibitor) vs. normalized response – variable slope” in GraphPad Prism (version 10.2.3).

### *Gaussia* Luciferase Assay

*Gaussia* luciferase activity was assessed in the cell culture supernatant using the Pierce *Gaussia* Luciferase Flash Assay Kit (Thermo Scientific) according to the manufacturer’s instructions. Cells were infected with recombinant VACVs encoding *Gaussia* luciferase under early (vEGluc), intermediate (vIGluc), or late (vLGluc) promoters. At indicated time points post-infection, supernatants were collected, and luminescence was recorded using a GloMax luminometer (Promega).

### Quantitative Real-Time PCR

Total DNA was extracted using the EZNA Blood DNA Kit (Omega Bio-Tek) according to the manufacturer’s protocol. Quantitative PCR was performed using the CFX96 Touch Real-Time PCR System (Bio-Rad) and All-in-One 2× qPCR mix (GeneCopoeia). VACV DNA was quantified using primers targeting the C11R gene and normalized to 18S rRNA levels as an internal control. The sequences for the primers used are as follows:

C11p FW: AAACACACACTGAGAAACAGCATAAA

C11p Rev: ACTATCGGCGAATGATCTGATTATC

18S rRNA FW: CGA TGC TCT TAG CTG AGT GT

18S rRNA Rev: GGT CCA AGA ATT TCA CCT CT

### *Ex Vivo* Mouse Lung Tissue Infection Assay

Lung tissues were harvested from C57BL/6 (B6) mice immediately after euthanasia. The lungs were rinsed in phosphate-buffered saline (PBS) and sliced into thin sections using a McIlwain tissue chopper. Equal weights of lung slices (4-5 slices) were distributed into each well of a 24-well tissue culture plate. Tissues were infected with 1 × 10⁵ PFU of vLGluc VACV in the presence of either 10 µM CPX or 40 µg/mL AraC or vehicle. At 1 hpi, the inoculum was removed and replaced with fresh media containing the respective compounds. *Gaussia* luciferase activity was measured directly from the culture supernatants at the indicated time points.

Experiments were performed using three independent biological replicates. All mouse experiments were conducted in accordance with protocols approved by the Texas A&M University Institutional Biosafety Committee (Approval of Animal Use Protocol IACUC 2022-0224).

### FerroOrange Assay for Intracellular Iron Detection

Approximately 30,000 HFFs were seeded in black-walled, clear-bottom 96-well plates and incubated overnight at 37 °C in a humidified 5% CO₂ incubator. The following day, cells were either mock-infected or infected with VACV in the presence of vehicle control or indicated compounds. At indicated times post-infection, cells were washed three times with Hank’s Balanced Salt Solution (HBSS), and 1 µmol/L FerroOrange (Cell Signaling Technology) in HBSS was added to each well. Plates were incubated at 37 °C for 30 minutes, and fluorescence was measured using a Cytation 5 imaging reader with Ex/Em settings of 543/580 nm.

### Transfection with plasmids expressing human RRM2 or VACV-F4L

The following mammalian expression plasmids were designed and obtained from VectorBuilder: Codon optimized VACV-F4L, human RRM2, or empty vector. The 293-FT cells were seeded in 24-well plates and transfected with 1 µg of the indicated plasmids using Lipofectamine 3000 reagent according to the manufacturer’s protocol. Six hours post-transfection (hpt), the culture medium was replaced with fresh complete DMEM. At 24 hpt, cells were infected with vLGluc VACV at an MOI of 0.01 in the presence of vehicle or 10 µM CPX. To preserve cell viability, the medium was not replaced after the 1-hour adsorption period. *Gaussia* luciferase activity was measured from the culture supernatant at 24 hours post-infection using a luminometer as described before.

### Statistical Analysis

Data are presented as mean ± standard deviation (SD) from at least three independent biological replicates, unless otherwise stated. Statistical analyses were performed using GraphPad Prism (version 10.2.3) and Microsoft Excel (version 16.43). For comparisons between two groups, a two-tailed paired t-test was used. For comparisons among more than two groups, one-way ANOVA was followed by Tukey’s or Dunnett’s post hoc tests. For two-factor comparisons, two-way ANOVA followed by Tukey’s post hoc test was applied. Statistical significance was defined as follows: ns, P > 0.05; *P ≤ 0.05; **P ≤ 0.01; ***P ≤ 0.001; ****P ≤ 0.0001.

## Results

### Ciclopirox efficiently suppresses VACV replication without cytotoxicity in HFFs

Ciclopirox (CPX, **Fig. 1A**) was identified as a top hit in suppressing VACV replication in our previous high-throughput screening of FDA-approved and bioactive compounds [34]. To validate its antiviral activity, we infected primary human foreskin fibroblasts (HFFs) with VACV at both high (MOI = 2) and low (MOI = 0.01) multiplicities of infection in the presence of 10 µM CPX or 40 µg/mL cytarabine (AraC). Plaque assay, which is the gold standard method to determine infectious viral yield [38], showed that CPX significantly reduced VACV titers compared to vehicle (DMSO)-treated controls by more than 1,000 folds (**Fig. 1B, C**), with suppression levels comparable to AraC at both MOIs. Neither compound significantly impacted cell viability as measured by trypan blue exclusion (**Fig. 1D**) indicating that the reduction in VACV replication was not due to cytotoxicity.

**Figure 1.**
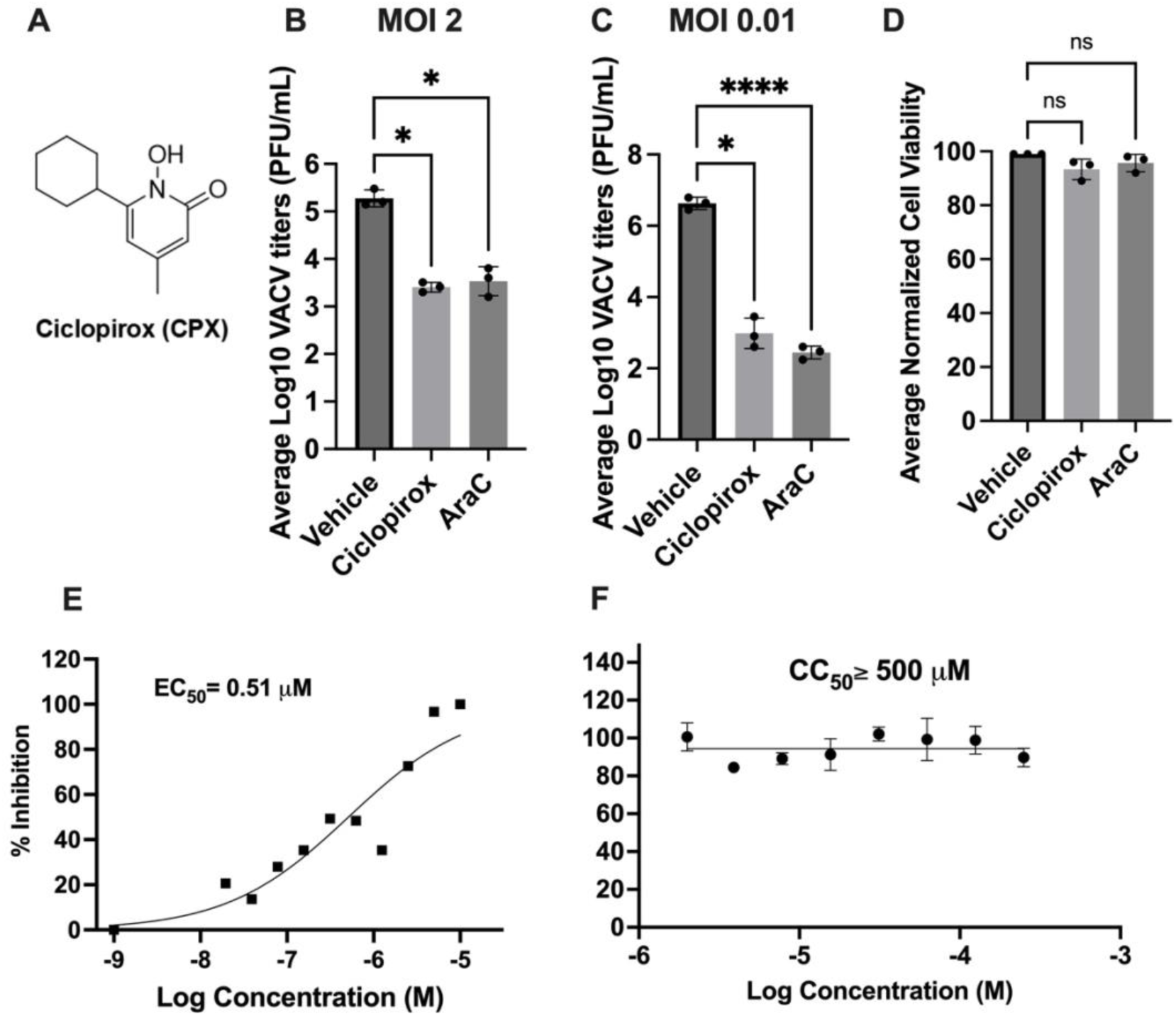
CPX inhibits VACV replication without affecting host cell viability. (A) Structures of ciclopirox (CPX). (B–C) HFFs were infected with VACV at MOI = 2 (B) or MOI = 0.01 (C) in the presence of 10 µM CPX, 40 µg/mL cytarabine (AraC), or vehicle (DMSO). Viral titers were quantified by plaque assay at 24 hpi for MOI 2 (B) or 48 hpi for MOI 0.01 (C). (D) Cell viability was assessed by trypan blue exclusion following treatment of HFFs with 10 µM CPX, 40 µg/mL cytarabine (AraC), or vehicle (DMSO) for 48 hours. (E) HFFs were infected with MOI = 0.01 of vLGluc in the presence of vehicle or a series of concentration of CPX. *Gaussia* luciferase activities were measured at 24 hpi to determine the EC_50_ value. (F) HFFs were treated with vehicle or a series of concentration of CPX. MTT assay was performed at 24 h post treatment to calculate the CC_50_ value. All data represent mean ± SD of at least three independent biological replicates. Statistical analysis was performed using one-way ANOVA with Dunnett’s post hoc test. *P ≤ 0.05; ****P ≤ 0.0001; ns, not significant.

To determine the potency of CPX, we next calculated the EC₅₀ using recombinant VACV expressing *Gaussia* luciferase under a late promoter (vLGluc). This assay system has been extensively optimized and validated for the assessment of antiviral activity and EC₅₀ determination for VACV inhibitors [34,39,40]. Using this approach, CPX exhibited a dose-dependent inhibition of VACV replication with an EC₅₀ of 0.51 µM (**Fig. 1E**). In parallel experiments, we previously determined that the EC₅₀ of brincidofovir against VACV is in the low micromolar range, while tecovirimat exhibited an EC₅₀ of approximately 10 nM under identical conditions [39,40]. Importantly, CPX exhibited minimal cytotoxicity, with a CC₅₀ greater than 500 µM over 24-48 h incubation (**Fig. 1F**). Together, these findings establish CPX as a potent and selective inhibitor of VACV replication in primary HFFs.

### CPX treatment suppresses VACV DNA replication and post-replicative gene expression

Poxvirus replication occurs in a well-defined sequence of events, beginning with viral entry and early gene expression, followed by uncoating, DNA replication, intermediate and late gene expression, and concluding with post-replicative processes such as virion morphogenesis, assembly, and cell-to-cell spread [41,42]. To identify the stage of the VACV life cycle affected by CPX, we utilized recombinant VACV reporters expressing *Gaussia* luciferase under early (vEGluc), intermediate (vIGluc), or late (vLGluc) viral promoters. HFFs infected with vEGluc showed no difference in luciferase activity upon treatment with CPX or AraC compared to vehicle (**Fig. 2A**), suggesting that early gene expression is not affected. In contrast, both vIGluc and vLGluc activities were significantly reduced in cells treated with CPX, to an extent comparable with AraC (**Fig. 2B, C**) indicating CPX treatment suppresses VACV intermediate and late gene expression. To determine whether this defect in post-replicative gene expression was due to impaired viral DNA replication, we quantified VACV DNA levels by qPCR at 8 hpi. CPX treatment significantly reduced viral DNA accumulation relative to vehicle-treated controls, consistent with the effects observed with AraC (**Fig. 2D**). These findings indicate that CPX inhibits VACV DNA replication, leading to subsequent suppression of intermediate and late gene expression.

**Figure 2.**
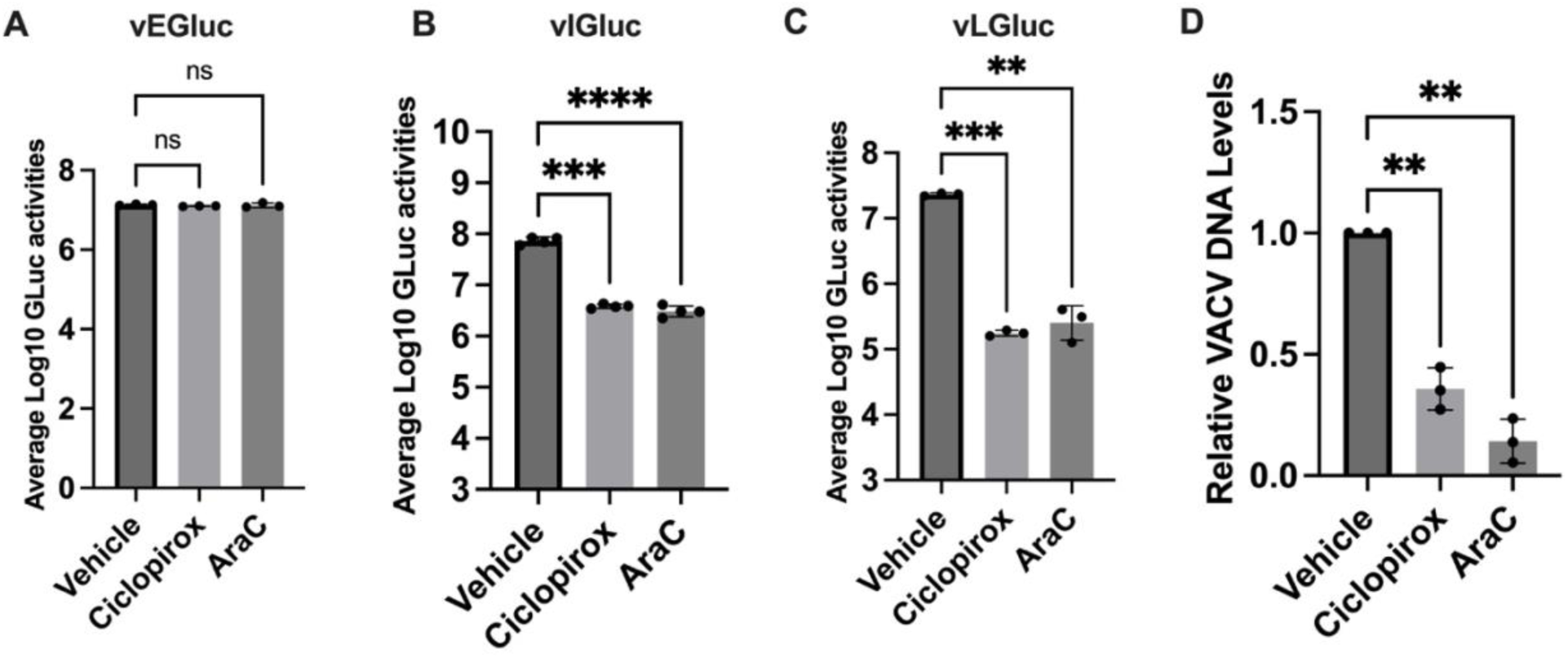
CPX suppresses VACV DNA replication and post-replicative gene expression. (A–C) HFFs were infected with recombinant VACV expressing *Gaussia* luciferase under early (vEGluc), intermediate (vIGluc), or late (vLGluc) promoters in the presence of vehicle, 10 µM CPX, or 40 µg/mL AraC. Luciferase activities in the supernatant were measured at 4 hpi for vEGluc and 8 hpi for vIGluc and vLGluc. (D) HFFs were infected with VACV at MOI = 2 in the presence of vehicle or 10 μM CPX for 8 h. Relative amounts of viral DNA were determined by real-time quantitative PCR using VACV-specific primers. Cells treated with 40 μg/mL AraC was used as a positive control. Statistical analysis was performed using one-way ANOVA with Dunnett’s post hoc test. **P ≤ 0.01; ***P ≤ 0.001; ****P ≤ 0.0001; ns, not significant.

### CPX suppresses VACV replication by interfering with iron-dependent processes

Given that CPX treatment led to a significant reduction in VACV DNA levels, we first tested whether this effect could be due to impaired nucleoside availability. We hypothesized that if CPX interfered with DNA replication by depleting nucleoside pools, supplementation with exogenous nucleosides would restore viral replication. We used nucleosides (AUGC) instead of nucleotides because nucleosides can readily enter cells and be phosphorylated to form nucleotides intracellularly. In contrast, nucleotides are negatively charged and do not easily cross cell membranes, making them ineffective for rescue experiments [43]. However, treatment with a combined mixture of adenosine, uridine, guanosine, and cytidine (AUGC) failed to rescue VACV replication at both high (MOI = 2) and low (MOI = 0.01) multiplicities of infection (**Fig. 3A, 3B**). These findings indicate that CPX’s antiviral effect is not due to impaired nucleoside availability or metabolism.

**Figure 3.**
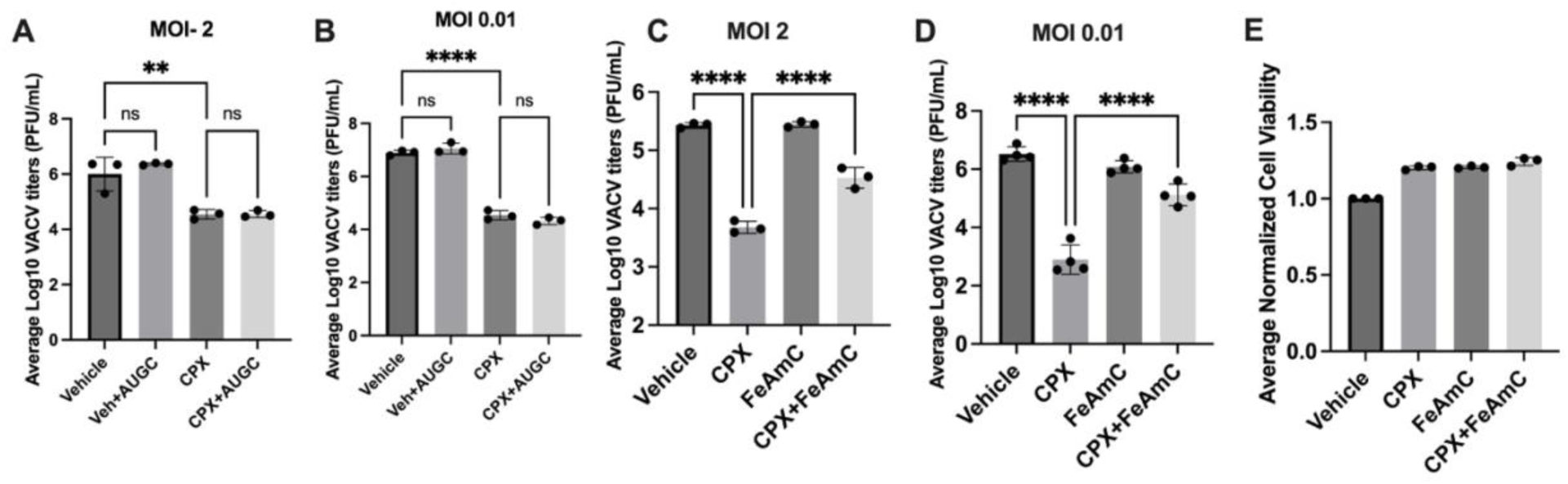
Effect of nucleoside and iron supplementation on rescuing VACV replication from CPX inhibition. (A, B) HFFs were infected with vLGluc VACV at MOI = 2 (A) or MOI = 0.01 (B). At 1 hpi, media was changed to vehicle or 10 μM CPX with or without 100 μM combined adenosine, uridine, guanosine, and cytidine (AUGC). VACV titers were measured at 24 hpi (MOI = 2) or 48 hpi (MOI = 0.01) by a plaque assay. (C) HFFs were infected with WT VACV (MOI = 2), and at 1 hpi, treated with vehicle, 10 μM CPX, 100 μM ferric ammonium citrate (FeAmC), or the combination of CPX + FeAmC. Virus titers were measured at 24 hpi by a plaque assay. (D) HFFs were infected with WT VACV (MOI = 0.01), and at 1 hpi, treated with vehicle, 10 μM CPX, 100 μM ferric ammonium citrate (FeAmC), or the combination of CPX + FeAmC. Virus titers were measured at 48 hpi by a plaque assay. (E) HFFs were treated with vehicle, 10 μM CPX, 100 μM FeAmC, or the combination of CPX + FeAmC. Cell viability was measured after 24 h of treatment using CCK8 assay. All values represent mean ± SD of at least three biological replicates. Statistical analysis was performed using one-way ANOVA with Dunnett’s post hoc test. *P ≤ 0.05; **P ≤ 0.01; ****P ≤ 0.0001; ns, not significant.

Ciclopirox is known to function as an iron chelator [44]. We therefore tested whether supplementation with ferric ammonium citrate (FeAmC) could restore VACV replication in the presence of CPX. FeAmC is a bioavailable source of Fe³⁺ that can enter cells and replenish intracellular iron levels [45,46], allowing us to test whether supplementing Fe³⁺ could counteract the iron-chelating effects of CPX on viral replication. Indeed, FeAmC co-treatment significantly restored VACV titers by more than 10 folds at MOI of 2 and more than 1000 folds at MOI of 0.01 in CPX-treated cells (**Fig. 3C, 3D**), supporting a role for iron metabolism in mediating CPX antiviral activity. Notably, cell viability remained unaffected across all treatment conditions, including CPX and FeAmC, suggesting that the observed antiviral effects were not due to cytotoxicity (**Fig. 3E**). Overall, these data suggest that CPX suppresses VACV replication not through disruption of nucleoside metabolism, but at least in part by interfering with iron-dependent processes.

### CPX treatment reduces intracellular iron levels, which is essential for VACV replication

Iron exists in multiple oxidation states and is compartmentalized both extracellularly and intracellularly where it supports critical metabolic and enzymatic processes [46,47], which may be required for efficient viral replication. To determine which iron pools are important for VACV, we tested chelators that selectively target different iron forms. Ciclopirox (CPX) and deferiprone (DFP) chelate intracellular Fe³⁺ [44,48–52], while deferoxamine (DFO) targets extracellular Fe³⁺ [44,51], and bathophenanthroline (BPhen) binds extracellular Fe²⁺[53,54].

VACV titers were significantly reduced by CPX and DFP at both high (MOI 2) and low (MOI 0.01) infection conditions (**Fig. 4A, 4B**), whereas BPhen and DFO did not significantly reduce VACV replication. This result suggests that intracellular Fe³⁺ is critical for VACV replication. In agreement with our findings, an earlier study has shown that BPhen treatment alone had no effect on VACV replication [54]. Previous studies have also shown that 2,2′-bipyridyl, an intracellular Fe²⁺ chelator, reduces VACV replication by targeting host ribonucleotide reductase activity [54], indicating this form of iron may also be important for VACV replication.

**Figure 4.**
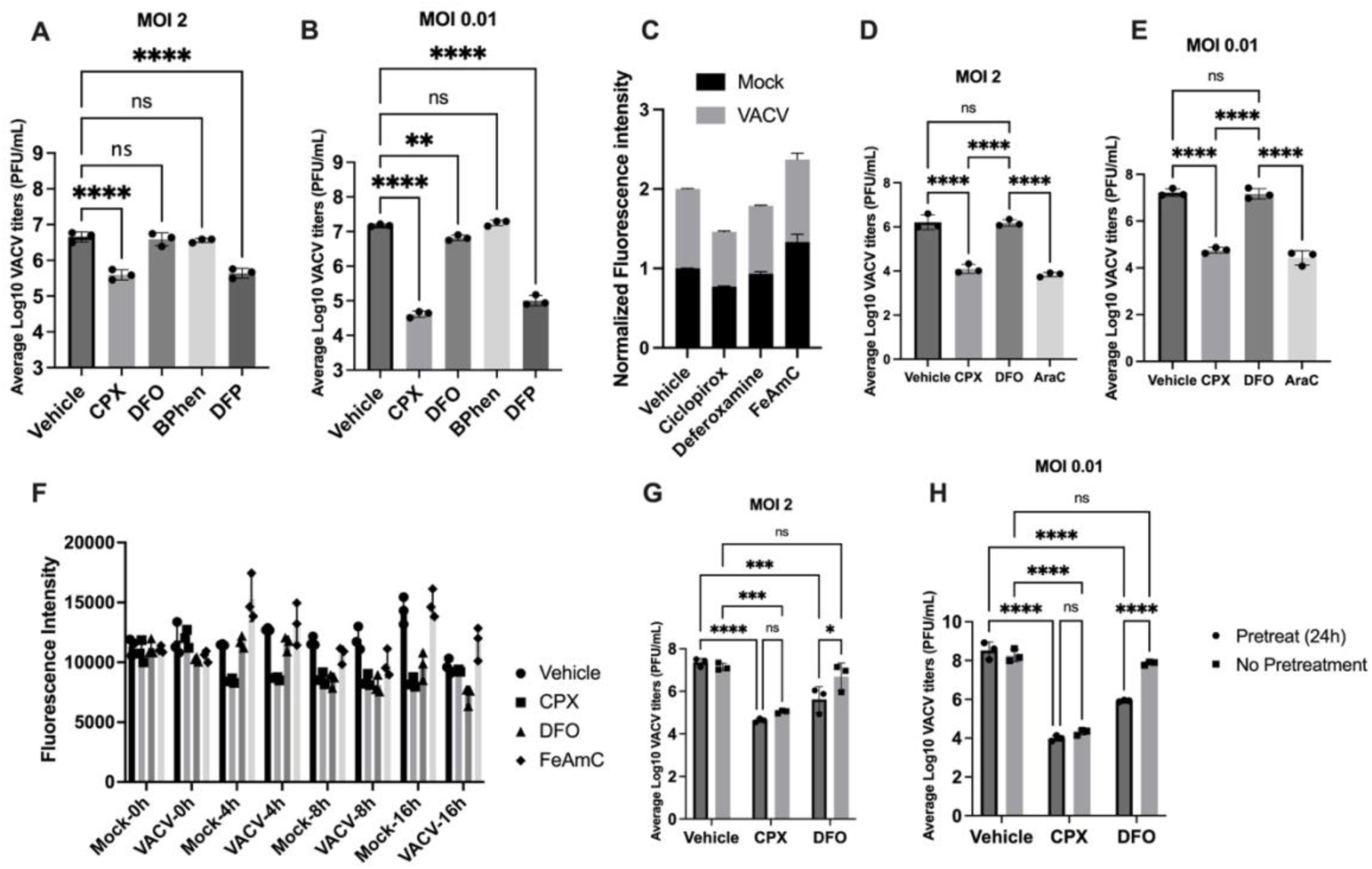
Intracellular iron is essential for VACV replication. (A–B) HFFs were infected with WT VACV at MOI = 2 (A) or MOI = 0.01 (B) and treated with vehicle, 10 μM CPX, 50 μM DFP, 50 μM DFO, or 50 μM bathophenanthroline (BPhen). Viral titers were measured at 24 hpi (MOI = 2) or 48 hpi (MOI = 0.01). (C) HFFs were infected with WT VACV at MOI = 2 and FerroOrange assay was performed at 4 hpi to measure intracellular Fe²⁺ levels in mock-or VACV-infected HFFs treated with vehicle, 10 μM CPX, 50 μM DFO, or 100 μM FeAmC. (D–E) HFFs were infected with WT VACV at MOI = 2 (D) or MOI = 0.01 (E) and treated with vehicle, CPX, or DFO at concentrations used above. Viral titers were measured at 24 hpi (MOI = 2) or 48 hpi (MOI = 0.01). (F) HFFs infected with MOI 2 of WT VACV in the presence of vehicle, CPX, or DFO. At indicated time points FerroOrange test was performed to measure intracellular Fe²⁺ depletion. All the decreases starting at 4hpi are statistically significant compared to vehicle treatment. (G–H) HFFs were pretreated with 10 μM CPX or 50 μM DFO for 24 h, then infected with VACV at MOI of 2 or 0.01 in the presence or absence of continued compound treatment. Viral titers were measured at 24 hpi for MOI = 2 (G) or 48 hpi for MOI = 0.01 (H). Data represent mean ± SD from at least three biological replicates. Statistical analysis was performed using one-way ANOVA with Dunnett’s post hoc test. *P ≤ 0.05; **P ≤ 0.01; ***P ≤ 0.001; ****P ≤ 0.0001; ns, not significant.

We next evaluated how CPX and DFO affected intracellular Fe²⁺ levels using FerroOrange, a fluorescent dye that selectively detects Fe²⁺. At 4 hpi, CPX significantly reduced FerroOrange signal in VACV-infected cells, while DFO had no measurable effect (**Fig. 4C**). Tools for specific detection of Fe³⁺ in live cells remain limited, so our focus was on Fe²⁺. To further probe the relevance of extracellular iron, we directly compared the antiviral activities of CPX and DFO across both MOIs. CPX consistently suppressed VACV replication, whereas DFO had minimal impact (**Fig. 4D, 4E**), supporting the idea that extracellular Fe³⁺ is less important for VACV replication.

A time-course analysis further revealed that CPX caused a rapid decrease in intracellular Fe²⁺ levels, starting at 4 hpi whereas DFO showed a slower response in reducing Fe²⁺ levels (**Fig. 4F**). These findings suggest that CPX acts directly on intracellular iron pools, while DFO’s effect is delayed and indirect due to its extracellular site of action.

Next, to determine whether early iron depletion alone is sufficient to inhibit infection, we pretreated cells with CPX or DFO for 24 h prior to infection, then continued or removed the treatment during infection. CPX pretreatment robustly suppressed VACV titers regardless of continued treatment (**Fig. 4G, 4H**), while DFO reduced VACV titers only when the pretreatment was followed by continued exposure during infection. These results indicate that CPX is capable of rapidly depleting critical intracellular iron pools to impair VACV replication, while DFO requires sustained treatment to achieve a similar effect. Overall, these findings demonstrate that VACV replication is highly dependent on intracellular iron availability.

### Overexpression of ribonucleotide reductase partially rescues CPX suppression of VACV replication

Next, we sought to determine whether the ability to rescue VACV replication from CPX inhibition was specific to iron or could be mediated by other metal ions. While iron supplementation with FeAmC successfully restored VACV replication, supplementation with FeCl₂, MgSO₄, or MnCl₂ failed to reverse CPX’s antiviral effect (**Fig. 5A, 5B**). This indicates that the rescue is iron-specific and not due to general metal ion supplementation. Although intracellular Fe²⁺ and Fe³⁺ are part of a redox-interchangeable labile iron pool [55], the inability of FeCl₂ to rescue viral replication further suggests that Fe²⁺ may be less important than Fe³⁺ in supporting VACV replication in the context of CPX treatment.

**Figure 5.**
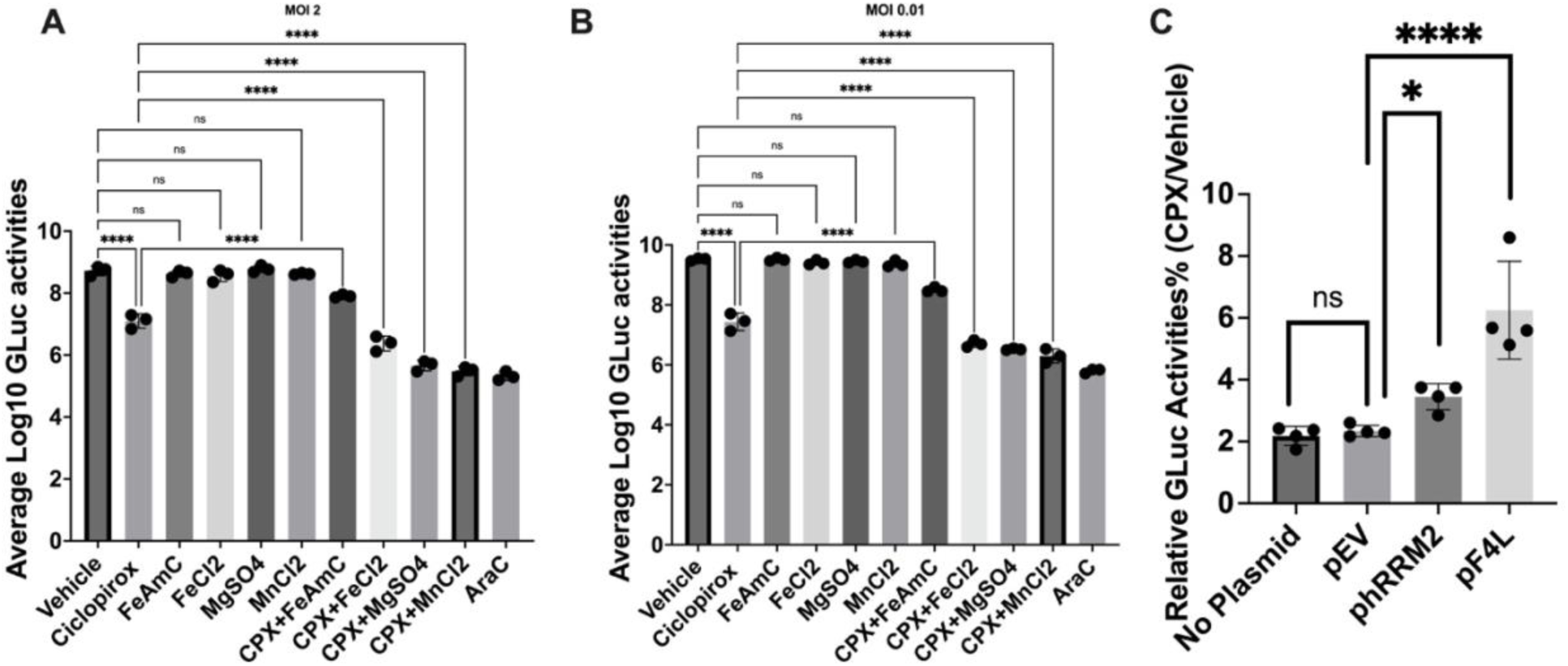
CPX suppresses VACV replication through disruption of iron-dependent ribonucleotide reductase activity. (A–B) HFFs were infected with VACV at an MOI of 2 (A) or MOI of 0.01 (B). At 1hpi, cells were treated with 10 µM CPX, 50 µM ferric ammonium citrate (FeAmC), 25 µM ferrous chloride (FeCl₂), 1 mM magnesium sulfate (MgSO₄), or 25 µM manganese chloride (MnCl₂), alone or in combination with CPX. Viral titers were measured at 24 hpi (A) or 48 hpi (B). (C) 293FT cells were transfected with plasmids encoding human RRM2, codon-optimized VACV F4L, or empty vector. At 24 h post-transfection, cells were infected with vLGluc VACV at MOI = 0.01 and treated with vehicle or 10 µM CPX. *Gaussia* luciferase activity in the supernatant was measured at 24 hpi. Data represent mean ± SD from at least three biological replicates. Statistical analysis was performed using one-way ANOVA with Dunnett’s post hoc test. *P ≤ 0.05; ****P ≤ 0.0001; ns, not significant.

Iron is a critical cofactor required for numerous biological processes, including the activity of ribonucleotide reductases (RNRs) that catalyze the conversion of ribonucleotides to deoxyribonucleotides, which are the building blocks essential for DNA replication [56,57]. Host cells express the iron-dependent RRM2 subunit of RNR [58,59], while VACV encodes a homologous viral RNR known as F4L [60,61]. F4L is essential for efficient VACV replication, particularly in normal cells, where RNR activity is limited compared to cancer cells [62,63]. The catalytic activity of both cellular RRM2 and the VACV-encoded F4L depends on the formation of a di-ferric tyrosyl radical cofactor, which is heavily dependent on Fe³⁺ [56,60,64]. This structural requirement underscores the importance of Fe³⁺ in supporting the function of these essential ribonucleotide reductases during viral DNA synthesis. Because supplementation with exogenous nucleosides (AUGC) failed to rescue VACV replication from CPX treatment (**Fig 3A, 3B**) and CPX treatment rapidly reduced intracellular iron concentrations (**Fig 4C, 4F**), we hypothesized that CPX may interfere with the activity of these iron-dependent enzymes. To test this, we transfected 293FT cells with plasmids containing either human RRM2 or a codon-optimized version of VACV F4L and infected them with VACV in the presence of CPX or absence of CPX. We observed that overexpression of either RRM2 or F4L led to significantly higher *Gaussia* luciferase activity compared to vector control under CPX treatment (**Fig. 5C**). These findings indicate that the antiviral effect of CPX is at least in part mediated by interference with the function of iron-dependent ribonucleotide reductases.

In our experiments, CPX and deferiprone, both of which are capable of chelating intracellular Fe³⁺, strongly suppressed VACV replication, while BPhen, which primarily targets extracellular Fe²⁺, had little to no effect. Our findings indicate that CPX disrupts intracellular iron homeostasis, as shown by rapid depletion in FerroOrange assays. However, the rescue by Fe³⁺ supplementation and failure of Fe²⁺ (FeCl₂) to restore viral replication from CPX suggest that bioavailable Fe³⁺ is functionally more important for VACV replication under CPX treatment.

This may be due to Fe³⁺’s role in replenishing the labile iron pool or stabilizing iron-dependent enzymes. Taken together, these results indicate that CPX inhibits VACV replication by interfering with intracellular iron availability, and that viral or cellular iron-dependent ribonucleotide reductases (F4L and RRM2) may be functionally impaired by CPX.

### CPX exhibits potent antiviral activity across multiple orthopoxviruses

Next, we tested whether CPX has broad-spectrum activity against other orthopoxviruses beyond VACV. To test this, we examined the effect of CPX treatment on the replication of CPXV, a zoonotic poxvirus that can elicit serious illness in humans and animals. HFFs infected with CPXV showed a significant reduction in viral titers upon treatment with 10 μM CPX compared to vehicle control (**Fig. 6A**). Next, we tested the ability of CPX to suppress the replication of MPXV-MA001 2022 isolate in primary HFFs. CPX treatment significantly suppressed MPXV replication at both high (MOI = 1) and low (MOI = 0.01) multiplicities of infection (**Fig. 6B–C**).

**Figure 6.**
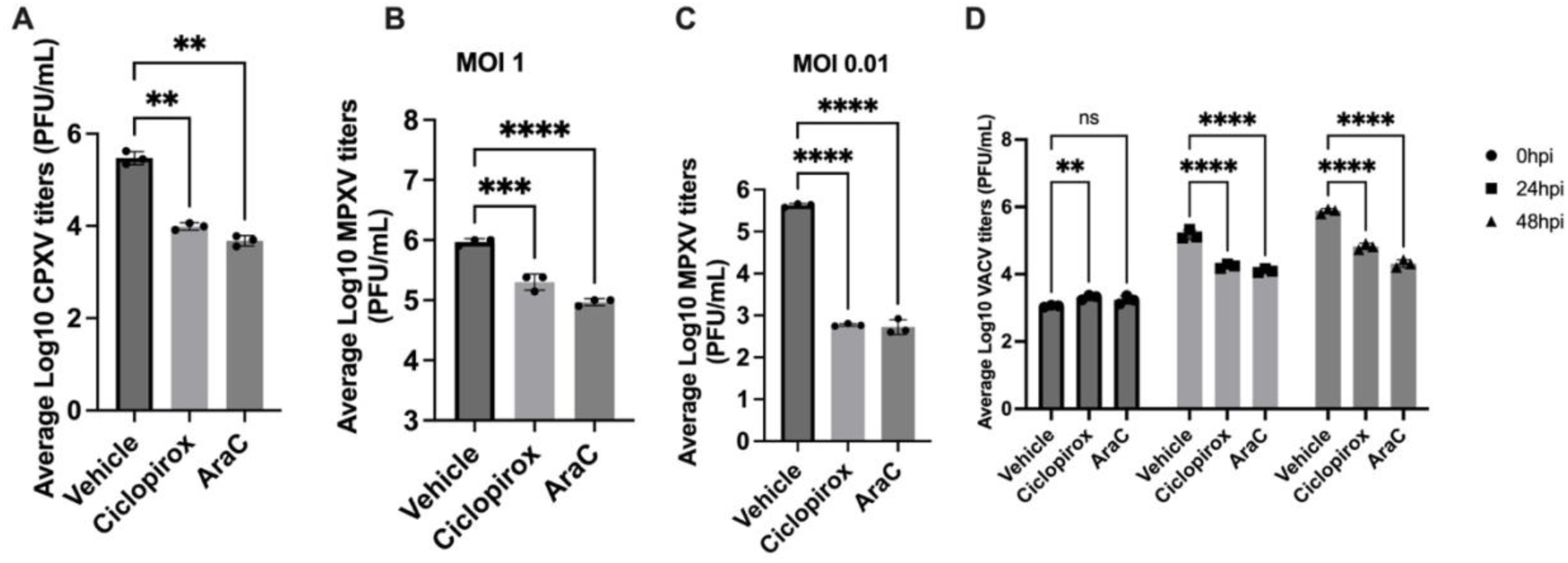
CPX inhibits replication of CPXV, MPXV, and VACV in HFFs and *ex vivo* mouse lung tissue. (A) HFFs were infected with CPXV at MOI = 0.01 and treated at 1 hpi with vehicle, 10 μM CPX, or 40 µg/mL AraC. Viral titers were measured at 48 hpi by a plaque assay. (B) HFFs were infected with MPXV at MOI =1 and treated at 1 hpi with vehicle, 10 μM CPX, or 40 µg/mL AraC. MPXV titers were measured at 24 hpi by a plaque assay. (C) HFFs were infected with MPXV at MOI = 0.01 and treated at 1 hpi with vehicle, 10 μM CPX, or 40 µg/mL AraC. MPXV titers were measured at 48 hpi by a plaque assay. (D) *Ex vivo* mouse lung tissues were infected with VACV and treated at 1 hpi with vehicle, 10 μM CPX, or 40 µg/mL AraC. VACV titers were measured at 0, 24, and 48 hpi by plaque assay. Each point represents an individual biological replicate. Error bars show standard deviation. Statistical significance was determined by one-way ANOVA with Tukey’s multiple comparisons test. **p < 0.01, ****p < 0.0001; ns = not significant.

To determine whether CPX can inhibit VACV replication in a more physiologically relevant setting, we infected *ex vivo* cultured mouse lung tissues with VACV and treated them with CPX, vehicle, or AraC. At both 24 and 48 hpi, CPX treatment robustly reduced viral titers in lung explants, similar to AraC, while vehicle treatment had no effect (**Fig. 6D**).

Together, these results indicate that CPX exhibits potent antiviral activity across multiple orthopoxviruses and remains effective in tissue-like environments, supporting its potential as a broad-spectrum antiviral candidate against orthopoxviruses.

## Discussion

This study identifies the FDA-approved antifungal agent CPX as a potent inhibitor of VACV replication by targeting host iron metabolism. CPX suppresses viral replication starting at the viral DNA replication stage by reducing intracellular iron levels. This disruption interferes with iron-dependent enzymes such as the host RRM2 and the viral ribonucleotide reductase homolog F4L. In addition to VACV, CPX’s antiviral activity extends to other orthopoxviruses, including CPXV and MPXV, highlighting its broader potential as a poxvirus inhibitor.

Our results demonstrate that CPX suppresses poxvirus replication through disruption of intracellular iron homeostasis, particularly by chelating Fe³⁺. The rescue of VACV replication by FeAmC, but not FeCl₂ or other divalent metal cations, suggests that bioavailable Fe³⁺ is especially important for VACV replication (Fig. 5A**–B**). Only intracellular Fe³⁺ chelators (CPX and DFP, which can access and chelate intracellular Fe³⁺, significantly suppressed VACV replication (Fig. 4A**–B**). Whereas DFP, which primarily chelates extracellular Fe³⁺ was largely ineffective when applied for short time periods (Fig. 4D**–E**), reinforcing the role of intracellular Fe³⁺. Prior studies have shown that the Fe²⁺ chelator 2,2′-bipyridyl reduces VACV replication by targeting RRM2 activity [54]. Although CPX treatment rapidly depletes intracellular Fe²⁺, as shown by FerroOrange test, this does not exclude the important role for Fe³⁺ in supporting VACV replication. FerroOrange specifically detects Fe²⁺ and does not account for changes in Fe³⁺ or total bioavailable iron. Given the dynamic redox interconversion between Fe²⁺ and Fe³⁺ in cells [55], the observed Fe²⁺ depletion likely reflects a broader iron-chelation effect.

Importantly, only Fe³⁺ supplementation (FeAmC), and not FeCl₂ or other divalent cations, rescued VACV replication from CPX treatment. Furthermore, only Fe³⁺-targeting chelators (CPX and DFP) robustly suppressed viral replication. Together, these findings suggest that although CPX reduces Fe²⁺ levels, the functional requirement for Fe³⁺, particularly for the activity of iron-dependent enzymes like RRM2 and F4L, is a more critical factor for viral DNA synthesis and replication in this context.

To investigate whether iron depletion alone could account for CPX’s antiviral effects, we examined the role of iron-dependent RNRs. VACV encodes F4L, a homolog of cellular RRM2, which is essential for DNA synthesis and replication [61,63]. Both host RRM2 and the VACV homolog F4L require a diferric-tyrosyl radical cofactor (Fe³⁺–Tyr) at their active sites to catalyze deoxyribonucleotide production [56,60,62,64]. This iron center is essential for initiating radical-based reduction of ribonucleotides, highlighting the role Fe³⁺ in their enzymatic function during DNA replication and supporting the role of Fe³⁺ in CPX mediated reduction of VACV titers.

Transient overexpression of either host RRM2 or VACV F4L partially attenuated the antiviral effects of CPX (Fig. 5C). Notably, F4L overexpression resulted in a more robust rescue compared to RRM2, suggesting that CPX may not solely act through a host-targeted mechanism but may also interfere directly with the viral enzyme F4L. This distinction is important, as many host-targeted antivirals could avoid resistance by sparing homologous viral proteins. Our findings raise the possibility that CPX may exert both host-directed and virus-directed effects.

These dual mechanisms of iron chelation impairing both host and viral RNR function highlights a previously underexplored vulnerability in poxvirus replication. In addition to RRM2, several cellular iron-sulfur cluster-containing DNA polymerases (e.g., Pol δ, Pol ε) and helicases (e.g., FANCJ, Dna2) rely on iron for proper folding, structural integrity, or enzymatic activity [65]. It is possible that additional VACV-encoded DNA replication factors also depend on iron for their functions. If CPX disrupts iron availability broadly within the cell, these additional replication and repair enzymes could also be functionally impaired. This broader impact on essential host replication machinery would further support CPX’s potential as a multi-target antiviral agent and underscore its therapeutic value against viruses that depend heavily on host DNA synthesis pathways. Another possibility is that CPX may interact with specific enzymes important for DNA replication, which can more specifically deplete iron to impair their enzymatic functions.

Ferroptosis has been linked to the replication of several viruses [66]. A recent report indicates that MPXV induces ferroptosis to facilitate viral replication and promotes inflammatory responses [67]. Ferroptosis is a form of regulated cell death driven by iron-dependent accumulation of lipid peroxides, particularly under conditions of oxidative stress. Intracellular Fe²⁺ promotes this process by catalyzing reactions that generate reactive oxygen species, leading to lipid membrane damage [68,69]. CPX, due to its strong iron-chelating properties, is generally reported to inhibit ferroptosis by limiting the availability of labile Fe²⁺ and thereby reducing lipid peroxidation [68,70]. Although VACV is shown to induce ferroptosis [67], our results suggest that this pathway is less likely to explain the antiviral effects of CPX for several reasons. First, CPX did not reduce the primary cell viability at the indicated concentrations for 24-48 h, where it selectively suppressed VACV replication (Fig. 1). Second, our results show that CPX effectively blocks VACV DNA replication (Fig. 2), which starts at an early time 2-3 hpi after VACV infection. Ferroptosis is less likely to explain the suppression of VACV DNA replication by CPX because it typically requires sustained oxidative stress and prolonged treatment, as it involves gradual accumulation of lipid peroxides and depletion of antioxidant defenses [68,69]. Third, Fe^2+^ is considered to play a major role in ferroptosis [71]. Our data indicate that Fe^3+^, but not Fe^2+^, can rescue VACV replication inhibited by CPX, suggesting additional iron-related mechanisms. Finally, our findings that iron supplementation partially rescues VACV replication and that overexpression of iron-dependent enzymes RRM2 and F4L attenuates CPX-mediated inhibition (Figs. 4–5) point toward targeted disruption of iron metabolism and enzyme function. While we do not rule out ferroptosis suppression may contribute to CPX inhibition of poxvirus replication at later stage of VACV infection after prolonged incubation, our observations support that CPX can suppress VACV replication at the early replication stage in a ferroptosis-independent mechanism, likely through impairing of iron-dependent enzymatic activity involved in viral DNA replication.

VACV is known to reprogram multiple aspects of host cell metabolism, including pathways involved in the TCA cycle, lipid biosynthesis, and nucleotide metabolism [72–74]. This study adds to the growing body of evidence by identifying a previously underappreciated role for iron metabolism in supporting VACV replication. Our findings also have implications for therapeutic development. CPX is an FDA-approved topical agent with a well-characterized safety profile, making it an attractive candidate for repurposing. Its broad activity against MPXV, CPXV (Fig. 6), and VACV, combined with a mechanism distinct from existing antivirals like tecovirimat and brincidofovir [6,7,75], suggests potential utility as a second-line or combination therapy. Importantly, CPX can affect both the host-dependent pathways and viral enzymes, which may limit the emergence of resistance, which is a critical advantage given the recent reports of tecovirimat resistance in MPXV [76].

While *ex vivo* mouse lung tissue provided physiologically relevant data (Fig. 6C), *in vivo* studies are needed to evaluate CPX efficacy, toxicity, and pharmacokinetics. Additionally, although our rescue and iron supplementation experiments implicate Fe³⁺dependent RNRs, direct measurement of RNR activity or binding between CPX and F4L or RRM2 are warranted to confirm these interactions. Ongoing efforts to improve the potency and safety profile of CPX will facilitate its potential repurposing for the treatment of MPXV and other poxvirus-related diseases.

Nevertheless, this work establishes CPX as a broad-spectrum orthopoxvirus inhibitor that acts primarily through intracellular iron depletion and interference with iron-dependent enzymes, including the viral ribonucleotide reductase F4L. These findings reveal an iron-dependent vulnerability in poxvirus replication and support further investigation of CPX and related iron-targeting agents as antiviral therapeutics.

## Funding sources

This work was supported in part by grants from the National Institutes of Health (R01AI183580) to Z.W. and Z.Y. The findings and conclusions in this report are those of the authors and do not necessarily represent the official position of the funding agencies.

## Declaration of competing interest

The authors declare that they have no known competing financial interests.

## Acknowledgements

We thank Nicholas Wallace (Kansas State University) for providing HFFs.

## Data availability

Data will be made available upon request by the corresponding author.

## Notes

### Competing Interest Statement

The authors have declared no competing interest.

## References

1. Rothenburg S, Yang Z, Beard P, Sawyer SL, Titanji B, Gonsalves G, et al. Monkeypox emergency: Urgent questions and perspectives. Cell. 2022;185: 3279–3281. doi:10.1016/j.cell.2022.08.002

2. Yang Z. Monkeypox: A potential global threat? J Med Virol. 2022;94: 4034–4036. doi:10.1002/jmv.27884

3. McCarthy M. Smallpox samples are found in FDA storage room in Maryland. BMJ. 2014;349: g4545. doi:10.1136/bmj.g4545

4. Noyce RS, Lederman S, Evans DH. Construction of an infectious horsepox virus vaccine from chemically synthesized DNA fragments. PLOS ONE. 2018;13: e0188453. doi:10.1371/journal.pone.0188453

5. Yang Z, Gray M, Winter L. Why do poxviruses still matter? Cell Biosci. 2021;11: 96. doi:10.1186/s13578-021-00610-8

6. Florescu DF, Keck MA. Development of CMX001 (Brincidofovir) for the treatment of serious diseases or conditions caused by dsDNA viruses. Expert Rev Anti Infect Ther. 2014;12: 1171–1178. doi:10.1586/14787210.2014.948847

7. Yang G, Pevear DC, Davies MH, Collett MS, Bailey T, Rippen S, et al. An orally bioavailable antipoxvirus compound (ST-246) inhibits extracellular virus formation and protects mice from lethal orthopoxvirus Challenge. J Virol. 2005;79: 13139–13149. doi:10.1128/JVI.79.20.13139-13149.2005

8. Adler H, Gould S, Hine P, Snell LB, Wong W, Houlihan CF, et al. Clinical features and management of human monkeypox: a retrospective observational study in the UK. Lancet Infect Dis. 2022;22: 1153–1162. doi:10.1016/S1473-3099(22)00228-6

9. Carvalho T. The unknown efficacy of tecovirimat against monkeypox. Nat Med. 2022;28: 2224–2225. doi:10.1038/d41591-022-00094-0

10. Friedberg DN. Hypotony and Visual Loss With Intravenous Cidofovir Treatment of Cytomegalovirus Retinitis. Arch Ophthalmol. 1997;115: 801–802. doi:10.1001/archopht.1997.01100150803021

11. Lea AP, Bryson HM. Cidofovir. Drugs. 1996;52: 225–230; discussion 231. doi:10.2165/00003495-199652020-00006

12. Vandercam B, Moreau M, Goffin E, Marot JC, Cosyns JP, Jadoul M. Cidofovir-induced end-stage renal failure. Clin Infect Dis Off Publ Infect Dis Soc Am. 1999;29: 948–949. doi:10.1086/520475

13. Erice A, Gil-Roda C, Pérez J-L, Balfour HH Jr, Sannerud KJ, Hanson MN, et al. Antiviral Susceptibilities and Analysis of UL97 and DNA Polymerase Sequences of Clinical Cytomegalovirus Isolates from Immunocompromised Patients. J Infect Dis. 1997;175: 1087–1092. doi:10.1086/516446

14. Chou S, Marousek G, Guentzel S, Follansbee SE, Poscher ME, Lalezari JP, et al. Evolution of mutations conferring multidrug resistance during prophylaxis and therapy for cytomegalovirus disease. J Infect Dis. 1997;176: 786–789. doi:10.1086/517302

15. Lurain NS, Chou S. Antiviral Drug Resistance of Human Cytomegalovirus. Clin Microbiol Rev. 2010;23: 689–712. doi:10.1128/cmr.00009-10

16. Desai AN, Thompson GR III, Neumeister SM, Arutyunova AM, Trigg K, Cohen SH. Compassionate Use of Tecovirimat for the Treatment of Monkeypox Infection. JAMA. 2022;328: 1348–1350. doi:10.1001/jama.2022.15336

17. The Antiviral Tecovirimat is Safe but Did Not Improve Clade I Mpox Resolution in Democratic Republic of the Congo | NIAID: National Institute of Allergy and Infectious Diseases. 14 Aug 2024 [cited 22 Jul 2025]. Available: https://www.niaid.nih.gov/news-events/antiviral-tecovirimat-safe-did-not-improve-clade-i-mpox-resolution-democratic-republic

18. Warner BM, Klassen L, Sloan A, Deschambault Y, Soule G, Banadyga L, et al. In vitro and in vivo efficacy of tecovirimat against a recently emerged 2022 monkeypox virus isolate. Sci Transl Med. 2022;14: eade7646. doi:10.1126/scitranslmed.ade7646

19. O’Laughlin K, Tobolowsky FA, Elmor R, Overton R, O’Connor SM, Damon IK, et al. Clinical Use of Tecovirimat (Tpoxx) for Treatment of Monkeypox Under an Investigational New Drug Protocol - United States, May-August 2022. MMWR Morb Mortal Wkly Rep. 2022;71: 1190–1195. doi:10.15585/mmwr.mm7137e1

20. NIH Study Finds Tecovirimat Was Safe but Did Not Improve Mpox Resolution or Pain | National Institutes of Health (NIH). [cited 22 Jul 2025]. Available: https://www.nih.gov/news-events/news-releases/nih-study-finds-tecovirimat-was-safe-did-not-improve-mpox-resolution-or-pain

21. Dittmar W, Lohaus G. HOE 296, a new antimycotic compound with a broad antimicrobial spectrum. Laboratory results. Arzneimittelforschung. 1973;23: 670–674.

22. Zangi M, Donald KA, Casals AG, Franson AD, Yu AJ, Marker EM, et al. Synthetic derivatives of the antifungal drug ciclopirox are active against herpes simplex virus 2. Eur J Med Chem. 2022;238: 114443. doi:10.1016/j.ejmech.2022.114443

23. Jue SG, Dawson GW, Brogden RN. Ciclopirox Olamine 1% Cream. Drugs. 1985;29: 330–341. doi:10.2165/00003495-198529040-00002

24. Sonthalia S, Agrawal M, Sehgal VN. Topical Ciclopirox Olamine 1%: Revisiting a Unique Antifungal. Indian Dermatol Online J. 2019;10: 481–485. doi:10.4103/idoj.IDOJ_29_19

25. Subissi A, Monti D, Togni G, Mailland F. Ciclopirox: recent nonclinical and clinical data relevant to its use as a topical antimycotic agent. Drugs. 2010;70: 2133–2152. doi:10.2165/11538110-000000000-00000

26. Inhibition of HIV-1 gene expression by Ciclopirox and Deferiprone, drugs that prevent hypusination of eukaryotic initiation factor 5A - ProQuest. [cited 21 Jul 2025]. Available: https://www.proquest.com/docview/1030132674?pq-origsite=gscholar&fromopenview=true&sourcetype=Scholarly%20Journals

27. Chen L, Chen D, Li J, He L, Chen T, Song D, et al. Ciclopirox drives growth arrest and autophagic cell death through STAT3 in gastric cancer cells. Cell Death Dis. 2022;13: 1007. doi:10.1038/s41419-022-05456-7

28. Bernier KM, Morrison LA. Antifungal drug ciclopirox olamine reduces HSV-1 replication and disease in mice. Antiviral Res. 2018;156: 102–106. doi:10.1016/j.antiviral.2018.06.010

29. Cáceres CJ, Angulo J, Contreras N, Pino K, Vera-Otarola J, López-Lastra M. Targeting deoxyhypusine hydroxylase activity impairs cap-independent translation initiation driven by the 5’untranslated region of the HIV-1, HTLV-1, and MMTV mRNAs. Antiviral Res. 2016;134: 192–206. doi:10.1016/j.antiviral.2016.09.006

30. Huang Z, Huang S. Reposition of the fungicide ciclopirox for cancer treatment. Recent Patents Anticancer Drug Discov. 2021;16: 122–135. doi:10.2174/1574892816666210211090845

31. Zhou H, Shen T, Luo Y, Liu L, Chen W, Xu B, et al. The antitumor activity of the fungicide ciclopirox. Int J Cancer J Int Cancer. 2010;127: 2467–2477. doi:10.1002/ijc.25255

32. Wei X, Zhou Y, Shen X, Fan L, Liu D, Gao X, et al. Ciclopirox inhibits SARS-CoV-2 replication by promoting the degradation of the nucleocapsid protein. Acta Pharm Sin B. 2024;14: 2505–2519. doi:10.1016/j.apsb.2024.03.009

33. Ciclopirox inhibits Hepatitis B Virus secretion by blocking capsid assembly | Nature Communications. [cited 21 Jul 2025]. Available: https://www.nature.com/articles/s41467-019-10200-5

34. Peng C, Zhou Y, Cao S, Pant A, Campos Guerrero ML, McDonald P, et al. Identification of Vaccinia Virus Inhibitors and Cellular Functions Necessary for Efficient Viral Replication by Screening Bioactives and FDA-Approved Drugs. Vaccines. 2020;8: 401. doi:10.3390/vaccines8030401

35. Pant A, Cao S, Yang Z. Asparagine Is a Critical Limiting Metabolite for Vaccinia Virus Protein Synthesis during Glutamine Deprivation. J Virol. 2019;93: e01834–18. doi:10.1128/JVI.01834-18

36. Cotter CA, Earl PL, Wyatt LS, Moss B. Preparation of Cell Cultures and Vaccinia Virus Stocks. Curr Protoc Microbiol. 2015;39: 14A.3.1-14A.318. doi:10.1002/9780471729259.mc14a03s39

37. Cao S, Realegeno S, Pant A, Satheshkumar PS, Yang Z. Suppression of Poxvirus Replication by Resveratrol. Front Microbiol. 2017;8. doi:10.3389/fmicb.2017.02196

38. Navarro-Forero S, Dsouza L, Yang Z. MVA titration by plaque assay using crystal violet staining in DF-1 cells. One Health Adv. 2023;1: 28. doi:10.1186/s44280-023-00031-x

39. Dsouza L, Pant A, Offei S, Priyamvada L, Pope B, Satheshkumar PS, et al. Antiviral activities of two nucleos(t)ide analogs against vaccinia, mpox, and cowpox viruses in primary human fibroblasts. Antiviral Res. 2023;216: 105651. doi:10.1016/j.antiviral.2023.105651

40. Pant A, Brahim Belhaouari D, Dsouza L, Priyamvada L, Wang Z, Navarro-Forero S, et al. Suppression of poxvirus replication by SC144. Antiviral Res. 2025;240: 106204. doi:10.1016/j.antiviral.2025.106204

41. Greseth MD, Traktman P. The Life Cycle of the Vaccinia Virus Genome. Annu Rev Virol. 2022;9: 239–259. doi:10.1146/annurev-virology-091919-104752

42. Moss B. Poxviridae: the viruses and their replication. 5th ed. In: Knipe DM, Howley PM, editors. Fields virology. 5th ed. Philadelphia, PA: Lippincott Williams & Wilkins; 2013. pp. 2129–2159.

43. Wright NJ, Lee S-Y. Toward a Molecular Basis of Cellular Nucleoside Transport in Humans. Chem Rev. 2021;121: 5336–5358. doi:10.1021/acs.chemrev.0c00644

44. Eberhard Y, McDermott SP, Wang X, Gronda M, Venugopal A, Wood TE, et al. Chelation of intracellular iron with the antifungal agent ciclopirox olamine induces cell death in leukemia and myeloma cells. Blood. 2009;114: 3064–3073. doi:10.1182/blood-2009-03-209965

45. Goralska M, Harned J, Fleisher LN, McGahan MC. The effect of ascorbic acid and ferric ammonium citrate on iron uptake and storage in lens epithelial cells. Exp Eye Res. 1998;66: 687–697. doi:10.1006/exer.1997.0466

46. Wang J, Pantopoulos K. Regulation of cellular iron metabolism. Biochem J. 2011;434: 365–381. doi:10.1042/BJ20101825

47. Roemhild K, von Maltzahn F, Weiskirchen R, Knüchel R, von Stillfried S, Lammers T. Iron metabolism: Pathophysiology and Pharmacology. Trends Pharmacol Sci. 2021;42: 640–656. doi:10.1016/j.tips.2021.05.001

48. Roberts DJ, Brunskill SJ, Doree C, Williams S, Howard J, Hyde CJ. Oral deferiprone for iron chelation in people with thalassaemia. Cochrane Database Syst Rev. 2007; CD004839. doi:10.1002/14651858.CD004839.pub2

49. Bohn M, Kraemer KT. Dermatopharmacology of ciclopirox nail lacquer topical solution 8% in the treatment of onychomycosis. J Am Acad Dermatol. 2000;43: S57–69. doi:10.1067/mjd.2000.109072

50. Sigle H-C, Thewes S, Niewerth M, Korting HC, Schäfer-Korting M, Hube B. Oxygen accessibility and iron levels are critical factors for the antifungal action of ciclopirox against Candida albicans. J Antimicrob Chemother. 2005;55: 663–673. doi:10.1093/jac/dki089

51. Wanner RM, Spielmann P, Stroka DM, Camenisch G, Camenisch I, Scheid A, et al. Epolones induce erythropoietin expression via hypoxia-inducible factor-1α activation. Blood. 2000;96: 1558–1565. doi:10.1182/blood.V96.4.1558

52. Victor Hoffbrand A. Deferiprone therapy for transfusional iron overload. Best Pract Res Clin Haematol. 2005;18: 299–317. doi:10.1016/j.beha.2004.08.026

53. Cowart RE, Singleton FL, Hind JS. A comparison of bathophenanthrolinedisulfonic acid and ferrozine as chelators of iron(II) in reduction reactions. Anal Biochem. 1993;211: 151–155. doi:10.1006/abio.1993.1246

54. Romeo AM, Christen L, Niles EG, Kosman DJ. Intracellular Chelation of Iron by Bipyridyl Inhibits DNA Virus Replication: RIBONUCLEOTIDE REDUCTASE MATURATION AS A PROBE OF INTRACELLULAR IRON POOLS *. J Biol Chem. 2001;276: 24301– 24308. doi:10.1074/jbc.M010806200

55. Kakhlon O, Cabantchik ZI. The labile iron pool: characterization, measurement, and participation in cellular processes(1). Free Radic Biol Med. 2002;33: 1037–1046. doi:10.1016/s0891-5849(02)01006-7

56. Li H, Stümpfig M, Zhang C, An X, Stubbe J, Lill R, et al. The diferric-tyrosyl radical cluster of ribonucleotide reductase and cytosolic iron-sulfur clusters have distinct and similar biogenesis requirements. J Biol Chem. 2017;292: 11445–11451. doi:10.1074/jbc.M117.786178

57. Zhang C, Liu G, Huang M. Ribonucleotide reductase metallocofactor: assembly, maintenance and inhibition. Front Biol. 2014;9: 104–113. doi:10.1007/s11515-014-1302-6

58. Reichard P. From RNA to DNA, Why So Many Ribonucleotide Reductases? Science. 1993;260: 1773–1777. doi:10.1126/science.8511586

59. Stubbe J, van der Donk WA. Ribonucleotide reductases: radical enzymes with suicidal tendencies. Chem Biol. 1995;2: 793–801. doi:10.1016/1074-5521(95)90084-5

60. Vaccinia virus-encoded ribonucleotide reductase: sequence conservation of the gene for the small subunit and its amplification in hydroxyurea-resistant mutants. [cited 24 Jul 2025]. doi:10.1128/jvi.62.2.519-527.1988

61. McCraith S, Holtzman T, Moss B, Fields S. Genome-wide analysis of vaccinia virus protein–protein interactions. Proc Natl Acad Sci. 2000;97: 4879–4884. doi:10.1073/pnas.080078197

62. Gammon DB, Gowrishankar B, Duraffour S, Andrei G, Upton C, Evans DH. Vaccinia Virus–Encoded Ribonucleotide Reductase Subunits Are Differentially Required for Replication and Pathogenesis. PLOS Pathog. 2010;6: e1000984. doi:10.1371/journal.ppat.1000984

63. Potts KG, Irwin CR, Favis NA, Pink DB, Vincent KM, Lewis JD, et al. Deletion of F4L (ribonucleotide reductase) in vaccinia virus produces a selective oncolytic virus and promotes anti-tumor immunity with superior safety in bladder cancer models. EMBO Mol Med. 2017;9: 638–654. doi:10.15252/emmm.201607296

64. Larsson A, Sjöberg BM. Identification of the stable free radical tyrosine residue in ribonucleotide reductase. EMBO J. 1986;5: 2037–2040. doi:10.1002/j.1460-2075.1986.tb04461.x

65. Zhang C. Essential functions of iron-requiring proteins in DNA replication, repair and cell cycle control. Protein Cell. 2014;5: 750–760. doi:10.1007/s13238-014-0083-7

66. Wang J, Zhu J, Ren S, Zhang Z, Niu K, Li H, et al. The role of ferroptosis in virus infections. Front Microbiol. 2023;14. doi:10.3389/fmicb.2023.1279655

67. Chuai X, Wang Y, Wang C, Ye T, Shen X, Zhou J, et al. Monkeypox virus induces ferroptosis to facilitate viral replication and promotes inflammatory responses. Emerg Microbes Infect. 2025 [cited 22 Jul 2025]. Available: https://www.tandfonline.com/doi/abs/10.1080/22221751.2025.2522877

68. Dixon SJ, Lemberg KM, Lamprecht MR, Skouta R, Zaitsev EM, Gleason CE, et al. Ferroptosis: An Iron-Dependent Form of Nonapoptotic Cell Death. Cell. 2012;149: 1060– 1072. doi:10.1016/j.cell.2012.03.042

69. Stockwell BR, Angeli JPF, Bayir H, Bush AI, Conrad M, Dixon S, et al. Ferroptosis: a regulated cell death nexus linking metabolism, redox biology, and disease. Cell. 2017;171: 273–285. doi:10.1016/j.cell.2017.09.021

70. Chen Z, Wang W, Abdul Razak SR, Han T, Ahmad NH, Li X. Ferroptosis as a potential target for cancer therapy. Cell Death Dis. 2023;14: 460. doi:10.1038/s41419-023-05930-w

71. Li J, Cao F, Yin H, Huang Z, Lin Z, Mao N, et al. Ferroptosis: past, present and future. Cell Death Dis. 2020;11: 88. doi:10.1038/s41419-020-2298-2

72. Pant A, Brahim Belhaouari D, Dsouza L, Yang Z. Stimulation of neutral lipid synthesis via viral growth factor signaling and ATP citrate lyase during vaccinia virus infection. J Virol. 2024;98: e0110324. doi:10.1128/jvi.01103-24

73. Dsouza L, Pant A, Pope B, Yang Z. Vaccinia growth factor-dependent modulation of the mTORC1-CAD axis upon nutrient restriction. J Virol. 2025;99: e0211024. doi:10.1128/jvi.02110-24

74. Pant A, Dsouza L, Cao S, Peng C, Yang Z. Viral growth factor- and STAT3 signaling-dependent elevation of the TCA cycle intermediate levels during vaccinia virus infection. PLOS Pathog. 2021;17: e1009303. doi:10.1371/journal.ppat.1009303

75. Grosenbach DW, Honeychurch K, Rose EA, Chinsangaram J, Frimm A, Maiti B, et al. Oral Tecovirimat for the Treatment of Smallpox. N Engl J Med. 2018;379: 44–53. doi:10.1056/NEJMoa1705688

76. Alarcón J, Kim M, Terashita D, Davar K, Garrigues JM, Guccione JP, et al. An Mpox-Related Death in the United States. N Engl J Med. 2023;388: 1246–1247. doi:10.1056/NEJMc2214921

